# Arena3D^web^: Interactive 3D visualization of multilayered networks supporting multiple directional information channels, clustering analysis and application integration

**DOI:** 10.1101/2022.10.01.510435

**Authors:** Maria Kokoli, Evangelos Karatzas, Fotis A. Baltoumas, Reinhard Schneider, Evangelos Pafilis, Savvas Paragkamian, Nadezhda T. Doncheva, Lars Juhl Jensen, Georgios A. Pavlopoulos

**Affiliations:** Institute for Fundamental Biomedical Research, BSRC “Alexander Fleming”, Vari, 16672, Greece; University of Luxembourg, Luxembourg Centre for Systems Biomedicine, Bioinformatics Core, Esch-sur-Alzette, Luxembourg; Institute of Marine Biology, Biotechnology and Aquaculture (IMBBC), Hellenic Centre for Marine Research (HCMR), Former U.S. Base of Gournes, Heraklion, 71003, Greece; Novo Nordisk Foundation Center for Protein Research, University of Copenhagen, Copenhagen N, DK- 2200, Denmark; Center of New Biotechnologies & Precision Medicine, Medical School, National and Kapodistrian University of Athens, Athens, 11527, Greece

**Author notes:** To whom correspondence should be addressed.; Correspondence may also be addressed to. The authors wish to be known that the first two authors should be regarded as joint First Authors. Present Address: Georgios A. Pavlopoulos, Institute for Fundamental Biomedical Research, BSRC “Alexander Fleming”, 34 Fleming Street, Vari, 16672, Greece.

## Abstract

Arena3D^web^ is an interactive web tool that visualizes multi-layered networks in 3D space. In this update, Arena3D^web^ supports directed networks as well as up to nine different types of connections between pairs of nodes with the use of Bézier curves. It comes with different color schemes (light/gray/dark mode), custom channel coloring, four node clustering algorithms which one can run on-the-fly, visualization in VR mode and predefined layer layouts (zig-zag, star and cube). This update also includes enhanced navigation controls (mouse orbit controls, layer dragging and layer/node selection), while its newly developed API allows integration with external applications as well as saving and loading of sessions in JSON format. Finally, a dedicated Cytoscape app has been developed, through which users can automatically send their 2D networks from Cytoscape to Arena3D^web^ for 3D multi-layer visualization. Arena3D^web^ is accessible at http://arena3d.pavlopouloslab.info or http://arena3d.org

## Introduction

Network biology has become a major domain in modern biology that captures associations among biomedical terms and entities such as organisms, diseases, genes, proteins, drugs and processes (1, 2). To be able to view these networks, many visualization tools have been proposed (3, 4). While Cytoscape (5, 6), Gephi (7), Tulip (8) and Pajek (9) are few of the most widely used applications, latest efforts focus on addressing more targeted problems. NORMA (10, 11) for example focuses on distinguishing annotated groups in a network using innovative layout strategies. OmicsNet (12) is developed to support multi-omics integration and network visual analytics. Graphia (13) comes with advanced layout algorithms to visualize a network in 3D, and STRING (14) uses an in-house graph visualizer to support multi-edged connections (different channels of information) between two nodes (proteins) (15).

Arena3D^web^ (16), the web-based successor to Arena3D (17, 18), is the first, fully interactive 3D web tool for processing, analyzing and visualizing multilayered networks. It allows advanced 3D visualizations to be created and navigated by providing a rich collection of layout algorithms, which one can apply on a selected set of layers individually or in combination, the ability to color layers, nodes and edges, and options to highlight and resize nodes based on topological features.

Here, we present an updated version of Arena3D^web^, which goes far beyond the aforementioned features. It now supports directed networks to allow visualization of regulatory networks, multi-edge connections to allow multiple types of edges between the same two nodes, and intra/inter-layer node clustering. The new version also features alternative color themes including a light mode suitable for designing figures for print, and enhanced navigation virtual reality (VR) functionality for viewing large networks in 3D. Finally, we provide import/export functions in JSON format and an Application Programming Interface (API) for integration with other applications such as connecting Cytoscape with Cytoscape with Arena3D^web^ through the new Arena3D Cytoscape app.

### UI/UX enhancement

In this version, Arena3D^web^ comes with major aesthetic improvements. The tool now offers three different color themes including light, gray and dark mode (Figure 1A). The light mode in particular enables users to produce publication-ready figures with a white background more easily, without the need to tweak the respective scene, layer and edge colors manually. In addition, several predefined layer layouts are also offered. To this end, initial 3D layer setups are produced automatically and include a zig-zag, a star and a cube layout (Figure 1B). In the star layout, a virtual sphere of 360 degrees is equally divided by the total number of layers whereas in the cube layout, each cube can contain up to 6 layers. In the case of more than 6 layers, additional cubes are created and placed next to each other.

**Figure 1.**
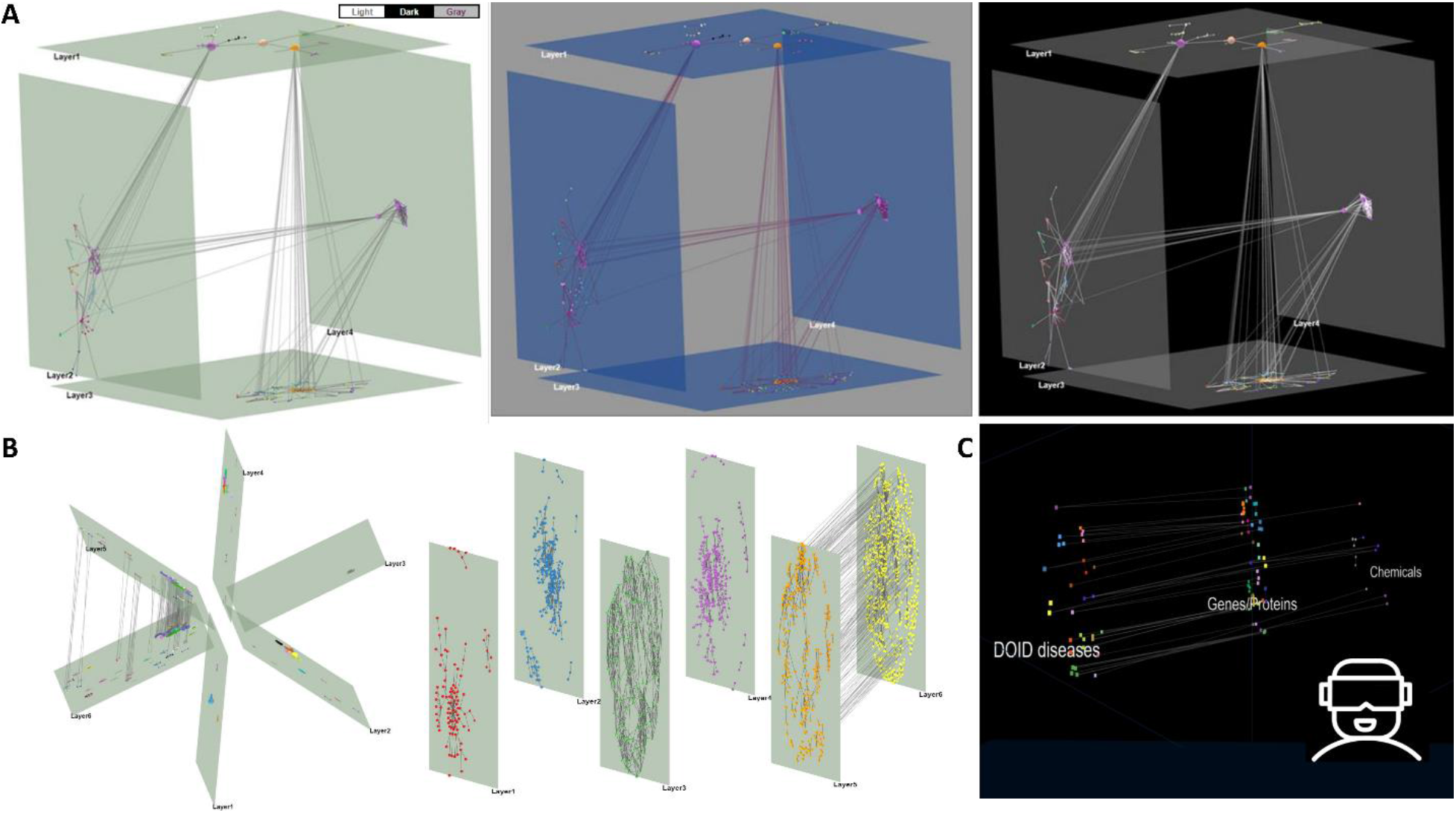
Updated viewing options. (A) Three default themes for a cube multi-layered network including (from left to right) a light, a gray and a dark mode option. (B) A star predefined layout (left) and a zig-zag layer layout (right) of six layers. (C) A VR view for a network with three layers (disease, proteins and chemicals).

In addition to the layouts, object manipulation has been significantly enhanced in this version. Raycasting via the *three*.*js* package has been implemented for easier layer and node selection with the mouse. Nodes change color on mouse hover as a visual cue while hovered layers are highlighted in red. Layer and node selection through double-click is also offered, while layers can be dragged with the mouse on the 3D scene as an alternative to the navigation control action buttons. The scene orbit controls (middle-click drag) have also been enhanced, allowing for smoother 3D scene rotations compared to the old version of the tool.

Finally, Arena3D^web^ now allows network exploration in VR mode (Figure 1C). VR views are static and can be accessed via a VR headset or a mobile phone with a gyroscope. For better clarity, layer floors are disabled while layer labels always face the user’s camera. Notably, the VR view is always offered in a new tab and visualizes the running view of Arena3D^web^ at any time point. For consistency, the objects’ coordinate system is always adjusted in the VR view. We note that the VR functionality is only accessible through the web application and not through the local version of Arena3D^web^, since all required VR files are served internally through the API (see section “Interoperability”).

## Multiple edge channels and directions

In biology, two entities can often be linked with more than one type of connection. In the STRING database, for example, two genes can be co-expressed, co-mentioned in literature or bind to each other. In a signal transduction network, a petri-net or a pathway, signal tracing as well as up- and down-stream analysis can be monitored by following directed paths visualized by arrows rather than plain lines. Therefore, the support of directed and multi-edged graphs is of great importance.

In this version of Arena3D^web^, we address both issues by allowing arrows and up to nine different types of connections between two nodes. In the case of multi-edged networks, each information channel must be labeled in the Arena3D^web^ input file and the corresponding edges are visualized as Bézier curves with distinct colors. In case of directed graphs, two intra- or inter-layer nodes can be connected via straight or curved arrows. The default channel colors are slightly adjusted to each theme but can be manually changed via color palettes at any time, while arrow sizes and curvature can be adjusted via dedicated sliders.

To demonstrate how Arena3D^web^ handles both directed and multi-edged connections, we provide two use cases. In the first case, we queried the STITCH (19) database for ‘*aspirin’* and filtered its interactions by only keeping the high-confidence ones (score > 0.7). We allow up to ten interactors for the channels: ‘*Experimental*’, ‘*Database*’ and ‘*Text Mining*’, depicted in green, blue and red, respectively (Figure 2A). Drug and protein nodes are separated into different layers. In the second example, we queried the PREGO knowledge base (20) for the ‘*anaerobic ammonium oxidation’* biological process (GO:0019331, *anammox*) and explored the associated microorganism taxa. The top 11 organisms (at the genus level) co-mentioned in the scientific literature with *anammox* were extracted (Figure 2B - left). For illustration purposes (Figure 2B - middle and right), the genus ‘*Beggiatoa’* was selected to show its associated environment types, including the top 12 environments from the *Literature* (text-mining) channel (edges indicated in green) along with the top 8 environments from the *Environmental Sample* channel (edges indicated in red).

**Figure 2.**
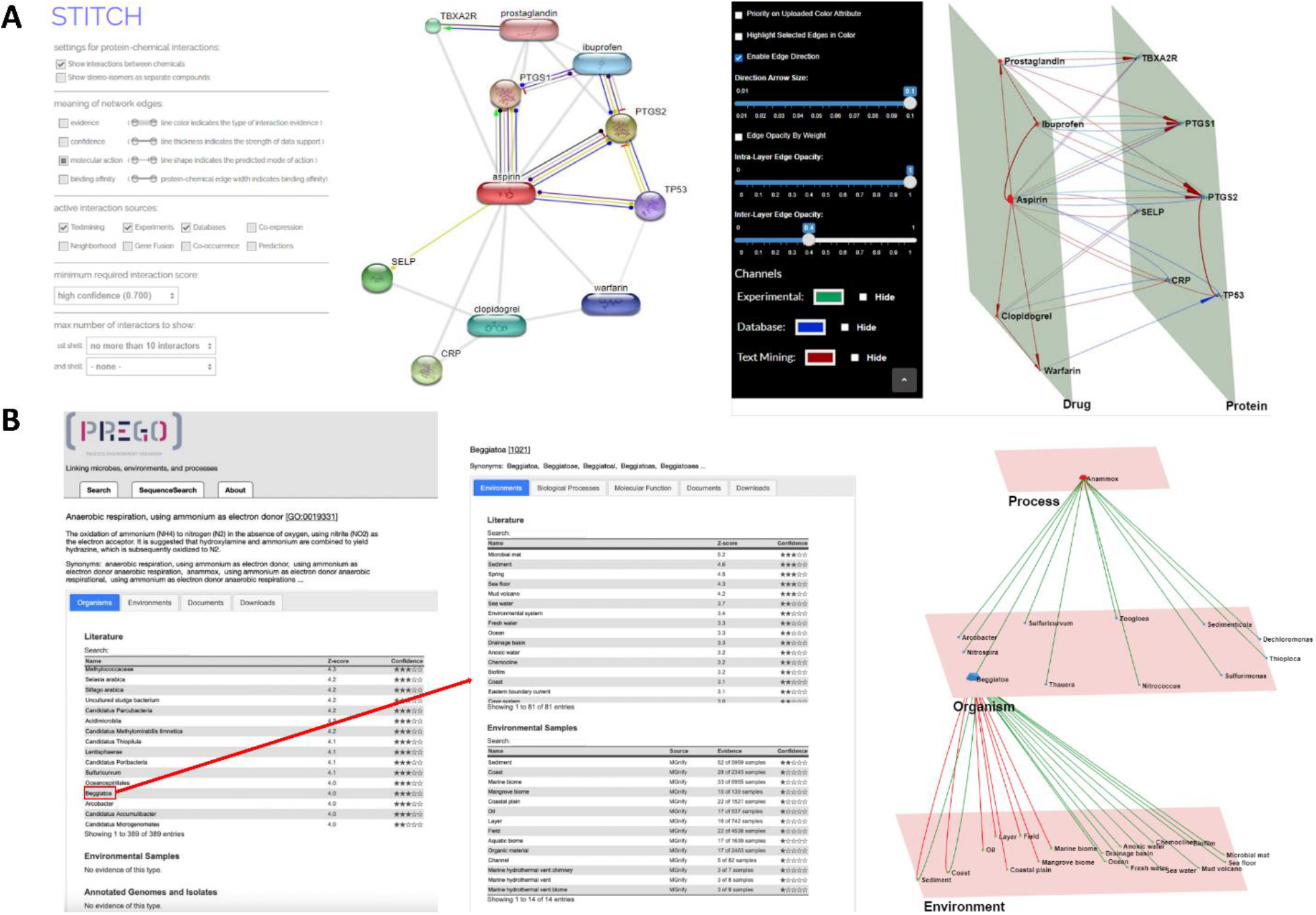
Multi-channel use cases. (A) A protein-chemical interaction network generated by STITCH. *Left:* Aspirin compound with its top ten interactors and molecular interaction channels is shown. *Right:* The same network in Arena3D^web^ format with three channels of information: “Experimental”, “Database” and “Text-Mining’’. Interactions are colored in green, blue and red edges respectively. The network consists of two layers, *Drugs* and *Proteins*. (?) PREGO’s *Process, Organism* and *Environment* association tables. *Left:* The associations of the ‘*anaerobic ammonium oxidation’* process with organisms. *Middle:* he associations of the genus *Beggiatoa* with the corresponding *Environments* and their respective channel sources (‘*Literature*’ (text-mining), ‘*Environmental Samples*’). *Right*: The multilayer and multi-edge Arena3D^web^ simultaneous visualization of these two distinct tables maintaining the source channel information (‘*Literature*’ (text-mining) shown via green edges, while ‘*Environmental Sample*’ derived-ones shown in red).

## Clustering

Arena3D^web^ now supports four main clustering algorithms offered by the *igraph* (21) library in combination with various global and local layouts. These are:

- *Louvain* (22): It is a multi-level modularity optimization algorithm for finding community structures and based on the modularity measure and a hierarchical approach.
- *Walktrap* (23): It performs random walks to detect densely connected neighborhoods. It is based on the assumption that random walks tend to restrict themselves in the same community.
- *Fast-Greedy* (24): It optimizes a modularity score to identify densely connected communities.
- *Label propagation* (25): It runs on a nearly linear time and tries to label the nodes with unique labels and update them by majority voting in the neighborhood of the node.

Users may apply any of these algorithms for a set of selected layers and channels to run them separately (per layer) or in combination (across layers). Upon clustering, the grouped nodes are placed on their respective layer according to the local layout selected from those offered by the *igraph* library. These layouts vary from simple ones like *circle, grid, star, random* to more advanced force-directed ones such as *Fruchterman–Reingold* (26), *Distributed Recursive (Graph) Layout* (27), *Multidimensional scaling* (28), *Kamada–Kawai* (29), *Large Graph Layout (LGL)* and *Graphopt* or hierarchical-based ones like *Reingold-Tilford* (30) and *Sugiyama*. A more detailed description of these algorithms can be found in our previous Arena3D^web^ article (16).

To minimize the cluster overlaps, Arena3D^web^ follows NORMA’s layout strategy named ‘Super nodes’ to visualize the clustered groups (11). To this end, a virtual super node is created to represent each cluster whereas the links among these super nodes (clusters) originate from initial node connections. After applying a global layout of preference, all of the initial nodes will be placed around their respective virtual super-nodes according to a user-selected local layout. Figure 3 demonstrates a visual example of the described procedure.

**Figure 3.**
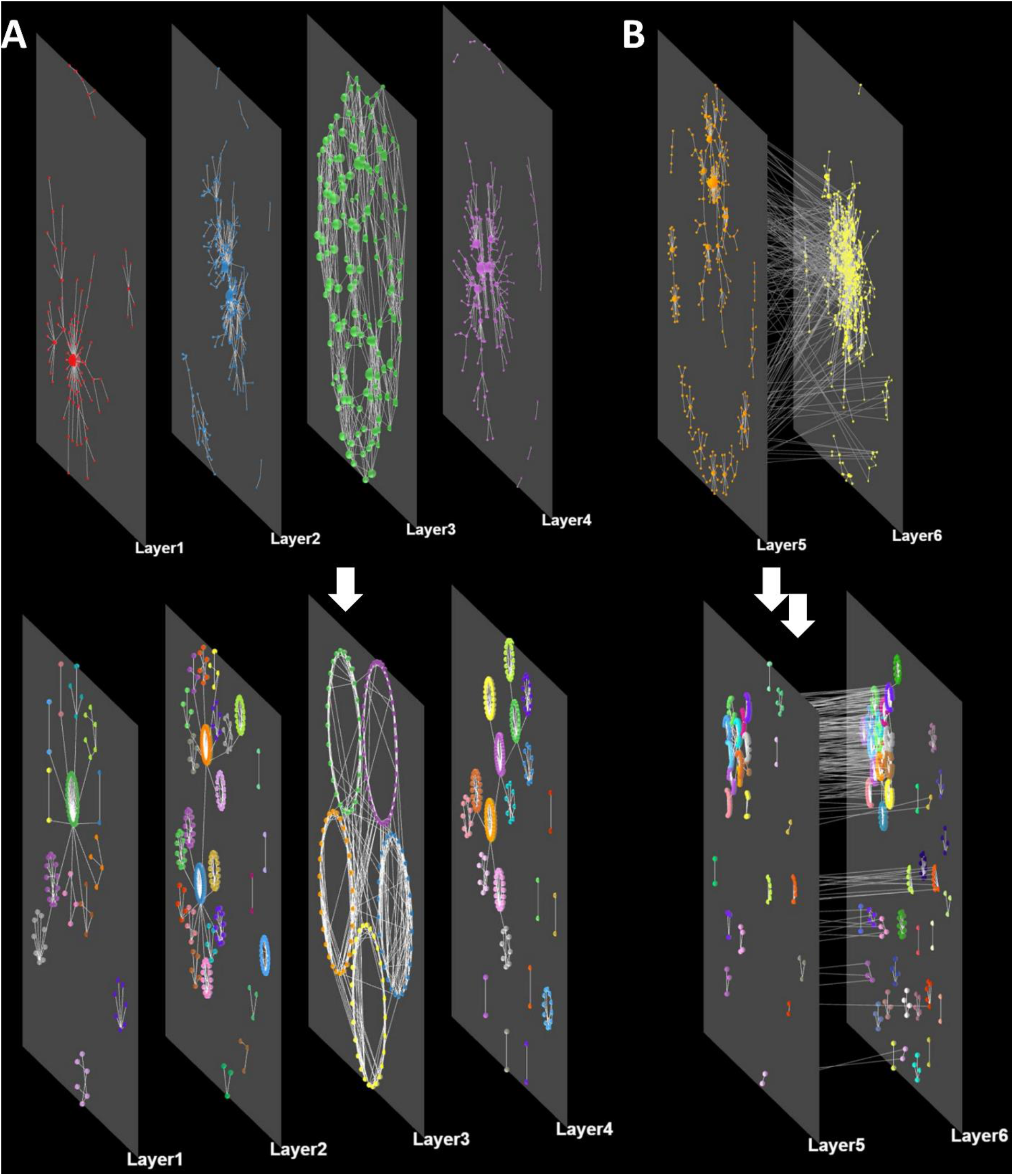
Layout and clustering combinations. (A) A network consisting of four different layers (no inter-layer edges) before and after applying the Louvain clustering algorithm per layer. A force-directed global layout and a circular local layout have been selected. (B) A network consisting of two interconnected layers before and after clustering with the same parameters as before.

## Integration with other tools

### Interoperability

In this version, Arena3D^web^ comes with its own API and a slightly updated input file format to allow multiple channels as well as a different import/export format for loading and saving sessions. Starting from the latter, Arena3D^web^ can save and load the running visualization in a JSON format. More specifically, the JSON object contains information regarding the: *(i)* scene object rotation, translation, scaling and background color, *(ii)* layer names, positions, rotations, scales, colors and widths, *(iii)* node colors, respective layers, positions, scales, colors and optional accompanying metadata, *(iv)* edge source and target nodes, opacities, colors, optional channels and *(v)*, an optional global label color as well as an optional flag for enabling edge direction.

The JSON format is suitable for calling Arena3D^web^ from external applications and services. For this purpose, a dedicated API route has been implemented in *Node*.*js*. This API call handles POST requests of an Arena3D^web^ JSON object, as described in the paragraph above (also see ?elp page, “API”) and opens the generated network object in an Arena3D^web^ viewer. For any missing parameters in the imported object, the tool provides default values where applicable.

To demonstrate the API’s usability, we provide two examples. Through the API utilization, Arena3D^web^ can be called from Darling (31), a text-mining disease-oriented software that produces 2D networks among tagged biomedical terms of ten different categories. In Darling, a user can query for diseases, chemicals, proteins, tissues or publications and then construct networks that link the relevant extracted biological entities based on literature co-mentions. In Arena3D^web^, the various biomedical entities will be placed on different layers according to their type (one layer per type). Similarly, through the recently published functional and literature enrichment analysis tool named Flame (32), two-layered Arena3D^web^ networks can be produced, one layer corresponding to the terms of a user-selected enriched category and a second layer containing the related genes/proteins. Two examples demonstrating the aforementioned APIs, are provided in Supplementary Figure 1.

**Supplementary Figure 1:**
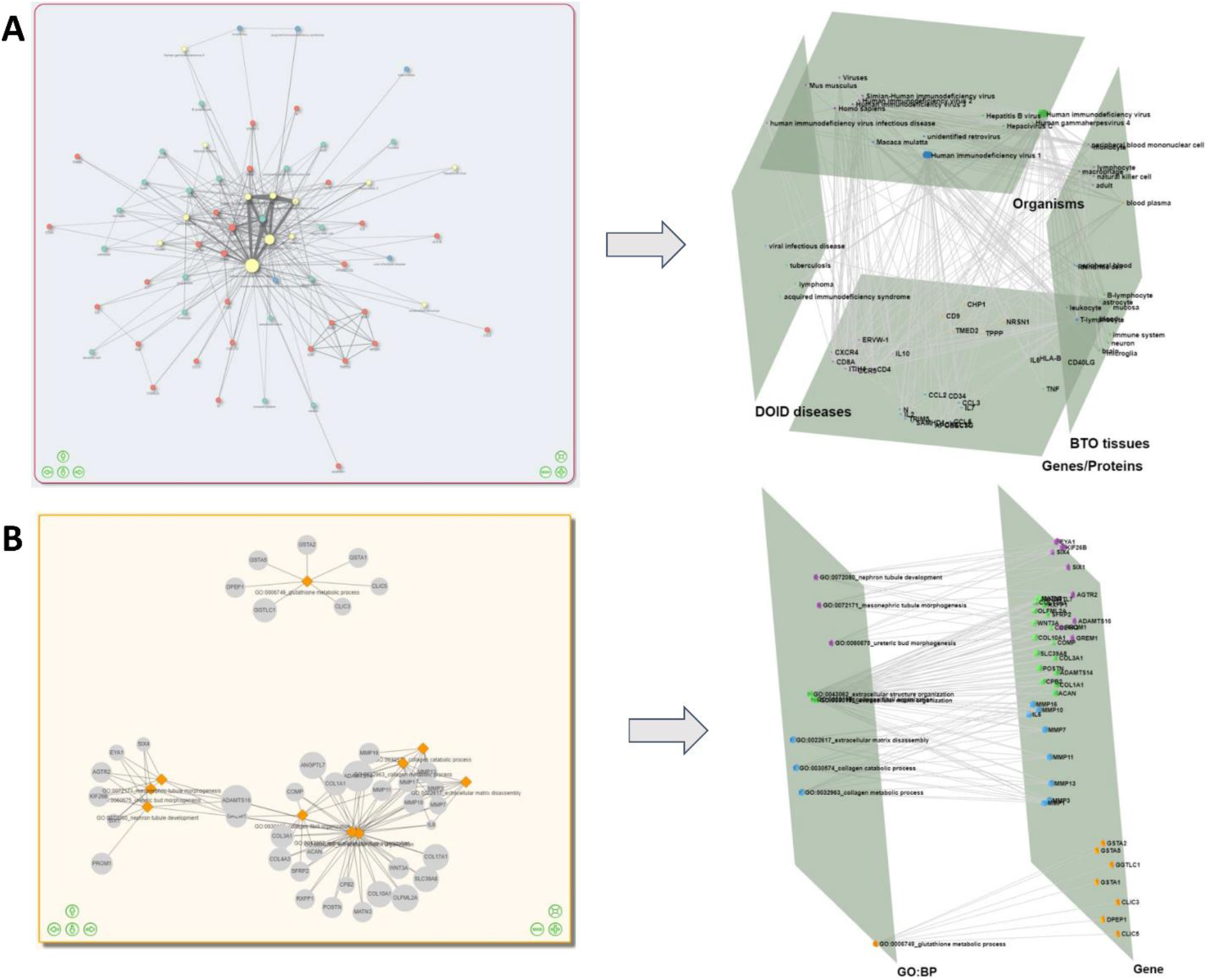
API functionality use cases. (A) A Darling query for HIV using the “Disease search” option to fetch literature from the DisGeNET dataset. 4 selected entity types were used (“DOID *Diseases*”, “*Genes/Proteins*”, “*BTO tissues*’’ and “*Organisms*”; see left). On the right, the returned biological entities are projected onto their respective layers in an Arena3D^web^ cube layout. (B) A Flame enriched network generated from a list of Idiopathic Pulmonary Fibrosis related differentially expressed genes and their top ten associated biological processes (see left). On the right, the genes and their related processes are projected in two distinct layers in Arena3D^web^.

### Integration with Cytoscape

Networks from Cytoscape can be sent to Arena3D^web^ through a dedicated Cytoscape app called *Arena3DwebApp*.. This app is implemented in Java and provides a simple interface, where the user can configure how the Cytoscape network will be transferred to Arena3D^web^. The most important setting is to choose which node attribute in Cytoscape should be used to define the layers; this can be any numeric or string value that defines up to 18 different non-overlapping groups. Arena3DwebApp retains much of the styling of the network from Cytoscape, extracting the currently displayed color, size and coordinates of each node as well as the directionality, color, thickness, and transparency of each edge. It also transfers the network-wide node label and network background colors. The user can choose which column to use for the node description and URL that can be seen in Arena3D^web^ as additional node information (on node right-click). If there are nodes that do not participate in any named layer, they are added to a layer named “*unassigned*” by default, but the user can choose to not import them in Arena3D^web^. The app generates a JSON file that is automatically sent to Arena3D^web^ and gets displayed in the user’s default web browser. If users want to share the layered network or open it later, they can export the JSON file from Cytoscape and import it in Arena3D^web^.

To illustrate the interoperability between Cytoscape and Arena3D^web^, we used stringApp v2.0 and Arena3DwebApp in combination from Cytoscape. Specifically, we used the “STITCH: protein/compound query” function of stringApp to search for the compound “aspirin” with a confidence score cutoff of 0.7 and up to ten additional interactors (compounds or human proteins). We then used stringApp to retrieve functional enrichment for the proteins interacting with aspirin and added all enriched diseases, tissues and KEGG pathways to the network (two, four, and five, respectively). The resulting network in Cytoscape is shown in Figure 4A. To transfer the network to Arena3D^web^, we opened the Arena3DwebApp dialog box shown in Figure 4B. We chose to use the column “stringdb::node type” to define the layers in Arena3D^web^, which means that chemical compounds will be placed in one layer, proteins in a second, and enriched terms in a third. We selected to not consider edges as directed, since STRING networks are undirected, and that the column “stringdb::description” should be used for node descriptions. Finally, we submitted this three-layer network to Arena3D^web^ for further 3D manipulations (Figure 4C). In the *enriched_term* layer, we show the three categories of enriched terms from STRING in three separate neighborhoods; KEGG pathways on top, tissues in the middle and diseases on the bottom.

**Figure 4:**
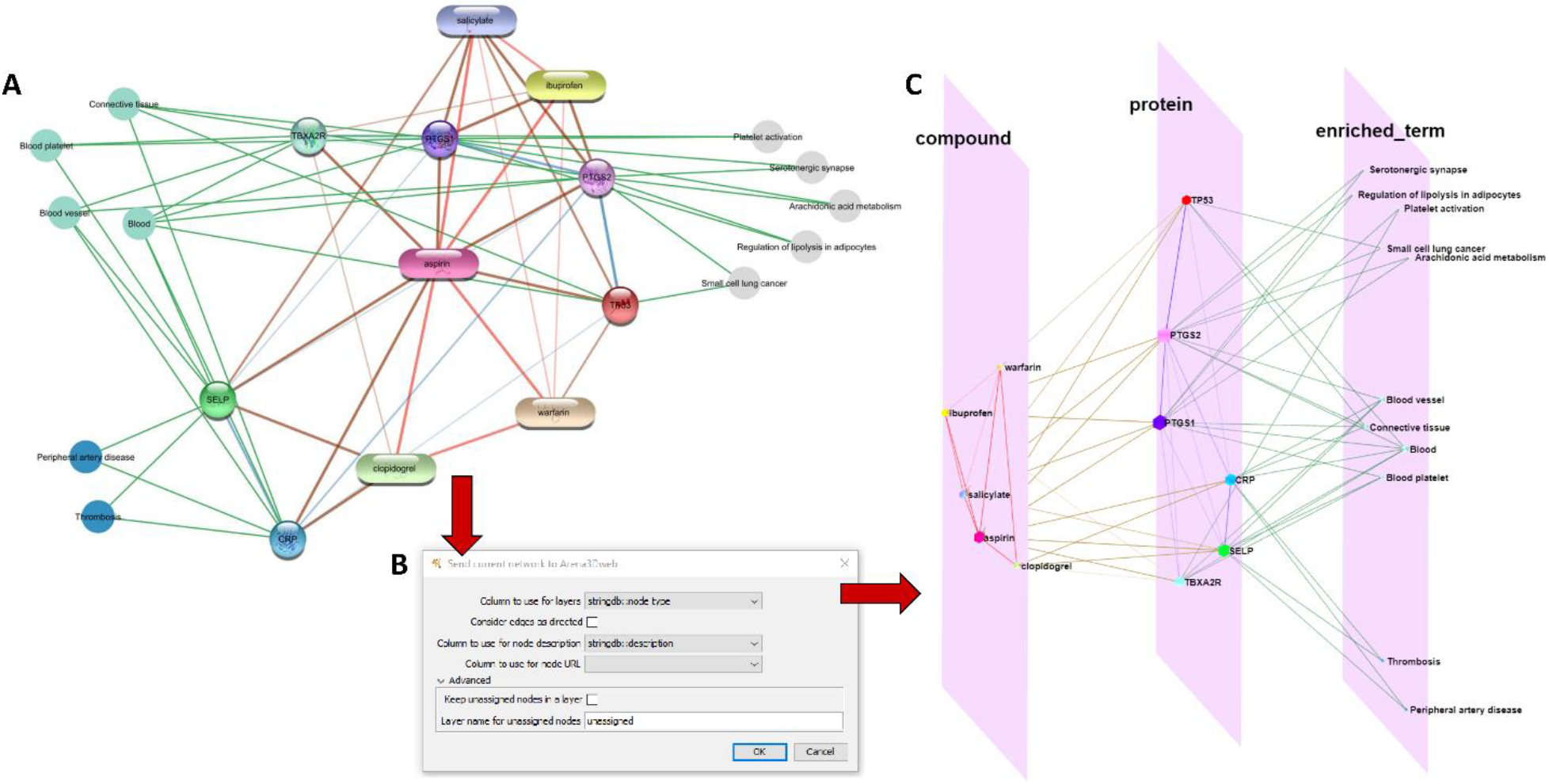
From Cytoscape to Arena3D^web^. (A) An aspirin chemical-protein interaction network along with enriched terms including diseases, tissues and KEGG pathways generated through stringApp in Cytoscape. Enriched disease nodes are colored red, tissues green and pathways gray. (B) The Arena3DwebApp dialogue window, where users are prompted to choose the node column that will indicate the various Layers in Arena3D^web^. Other options include edge directionality, columns for node descriptions and URLs and how to handle nodes without layer information. (C) The generated Arena3D^web^ network in three layers; compounds, proteins and enriched terms. Enriched pathways are placed on top, tissues in the middle and diseases on the bottom of the layer. The chemical compound interactions channel is colored red, compound-protein channel in brown, protein-protein in blue and protein participation in the various enriched terms in green. The node coloring scheme is in agreement with the initial STRING network in Cytoscape.

## SUMMARY

Arena3D^web^ is an innovative network analysis tool dedicated to the visualization of multilayered networks in 3D. Its main purpose is to visualize heterogeneous information of higher complexity in meaningful and appealing ways and simultaneously offer an interface for the production of high-quality, publication-ready and story-telling figures. While Arena3D^web^ retains an interdisciplinary character and is able to serve case-studies from various fields, future directions will include time series analyses and major integration with: *(i)* biomedical repositories for automated data extraction and information retrieval (e.g., PPI databases (33)) *(ii)* biomedical text-mining applications (e.g., STRING (14), BioTextQuest (34), STITCH (19), EXTRACT (35), OnTheFly (36), UniRed (37), PREGO (20)) and *(iii)* functional enrichment analysis tools (e.g., g:Profiler (38), aGOtool (39), DAVID (40), Panther (41)). To this end, integration with Cytoscape addresses many of these challenges, as several functionalities are offered through the available Cytoscape apps, whose results can be exported and visualized with Arena3D^web^ directly through the Arena3D^web^ Cytoscape app.

## AVAILABILITY

Arena3D^web^ is an open source application. Its code can be downloaded from the GitHub repository: https://github.com/PavlopoulosLab/Arena3Dweb.

The service is available online at http://arena3d.pavlopouloslab.info or http://arena3d.org. The API is accessible through https://bib.fleming.gr/bib/api/arena3dweb

The Arena3D^web^ Cytoscape plugin is available at: https://apps.cytoscape.org/apps/arena3DwebApp

